# Screening of Natural Plant Extracts for Antimicrobial Activity Against *Streptobacillus moniliformis*

**DOI:** 10.1101/2025.04.05.647341

**Authors:** Kishlay Kant Singh, Mansi Saini, Divya Prakash

## Abstract

The rise of antimicrobial resistance has driven the search for alternative antibacterial agents, including plant-based compounds. This study evaluates the antimicrobial potential of selected herbal extracts against *Streptobacillus moniliformis* using the agar well diffusion method. The tested extracts included Tulsi leaves (Ocimum sanctum), Neem leaves (Azadirachta indica), Bael leaves (Aegle marmelos), Adarak peels (Zingiber officinalef), Moringa seeds and leaves (Moringa oleifera), Dalchini (Cinnamomum verum), Lemon/Orange peels (Citrus limon and Citrus sinensis), and Ginger peels (Zingiber officinale). Among these, Bael leaves and Lemon peels demonstrated significant antibacterial activity, forming distinct zones of inhibition. In contrast, Neem and Moringa extracts did not inhibit bacterial growth. The observed antimicrobial activity is likely due to the presence of bioactive compounds such as flavonoids, tannins, and essential oils, which may disrupt bacterial cell walls and metabolic processes. Notably, *S. moniliformis* exhibited limited survival in culture, while other bacterial strains showed minimal resistance. These findings suggest that certain herbal extracts, particularly Bael leaves and Lemon peels, may serve as natural antimicrobial agents against *S. moniliformis*. Further studies are required to isolate and characterize the active compounds responsible for this antibacterial activity to explore their potential in developing alternative antimicrobial therapies.

## 1. Introduction

The increasing prevalence of antibiotic resistance has become a major global health concern, necessitating the search for alternative antimicrobial agents. Streptobacillus moniliformis, the causative agent of rat-bite fever, is a fastidious, gram-negative bacterium that can lead to systemic infections in humans [1]. Though it is not commonly associated with multidrug resistance, its ability to cause severe illness underscores the importance of identifying natural compounds with antibacterial potential. The use of plant-based antimicrobials, particularly herbal extracts, has gained significant attention due to their bioactive properties and historical use in traditional medicine [2].

Herbal extracts are derived from plants rich in biologically active compounds, such as flavonoids, alkaloids, tannins, and essential oils, which have been shown to possess antimicrobial, anti-inflammatory, and antioxidant properties [3]. These natural products offer a sustainable and environmentally friendly alternative to synthetic antibiotics, which often contribute to the emergence of resistant bacterial strains [4]. Furthermore, the diverse phytochemical compositions of herbal extracts can target multiple bacterial pathways, potentially reducing the risk of resistance development [5].

In this study, the antimicrobial activity of selected herbal extracts, including Tulsi leaves (Ocimum sanctum), Neem leaves (Azadirachta indica), Bael leaves (Aegle marmelos), Adarak peels (Zingiber officinale), Moringa seeds and leaves (Moringa oleifera), Dalchini (Cinnamomum verum), Lemon/Orange peels (Citrus limon and Citrus sinensis), and Ginger peels (Zingiber officinale), was evaluated against S. moniliformis [6]. These plants are known to contain bioactive compounds such as eugenol, tannins, flavonoids, cinnamaldehyde, and vitamin C, which may interfere with bacterial cell wall integrity, disrupt metabolic pathways, and inhibit quorum sensing [7].

Previous studies have demonstrated the antimicrobial potential of these herbal extracts against a wide range of bacterial pathogens, but their specific effects on S. moniliformis remain unexplored [8]. The primary objective of this study is to assess which extracts exhibit inhibitory effects on S. moniliformis and to investigate their potential mechanisms of action [9]. The findings from this research may contribute to the development of alternative antimicrobial therapies and highlight the importance of natural compounds in combating bacterial infections [10].

The agar well diffusion method was employed to evaluate the antibacterial activity of the selected extracts by measuring zones of inhibition around the wells. This widely used microbiological technique provides a clear indication of the efficacy of antimicrobial agents against bacterial growth [11].

This study aims to expand the scientific understanding of herbal extracts as natural antimicrobial agents against S. moniliformis, emphasizing their potential role in sustainable healthcare solutions [12]. By exploring plant-derived antimicrobials, this research addresses the ongoing challenge of bacterial infections and supports the growing demand for eco-friendly and effective therapeutic alternatives [13].

## 2. Materials & Methodology

Fresh samples of the following plants were collected from local sources: Tulsi leaves (Ocimum sanctum), Neem leaves (Azadirachta indica), Bael leaves (Aegle marmelos), Adarak peels (Zingiber officinale), Moringa seeds and leaves (Moringa oleifera), Dalchini (Cinnamomum verum), Lemon/Orange peels (Citrus limon and Citrus sinensis), and Ginger peels (Zingiber officinale) [14]. Each plant species was authenticated by a botanist to ensure accuracy. These plants were selected based on their reported antibacterial properties and historical use in traditional medicine [14].

The collected plant materials were processed for extraction and antimicrobial evaluation against Streptobacillus moniliformis [15]. The study aimed to determine the efficacy of these herbal extracts as potential natural alternatives to conventional antibiotics, addressing the need for alternative antimicrobial strategies [16].

### 2.1. Plant Material collection

The herbal extracts selected for this study included Tulsi leaves (*Ocimum sanctum*), Neem leaves (*Azadirachta indica*), Bael leaves (*Aegle marmelos*), Adarak peels (*Zingiber officinale*), Moringa seeds and leaves (*Moringa oleifera*), Dalchini (*Cinnamomum verum*), Lemon/Orange peels (*Citrus limon* and *Citrus sinensis*), and Ginger peels (*Zingiber officinale*), as listed in Table 1. These plants were chosen based on their well-documented antimicrobial properties and traditional medicinal usage [17].

**Table 1:** Catalog of herbal extracts used & herbal applications.

| S. No. | Herbs Name | Family Name | Part Used | Ayurvedic/Traditional Uses |
| --- | --- | --- | --- | --- |
| 1 | Neem (*Azadirachta indica*) | Meliaceae | Leaves | Antibacterial, Antifungal, Skin Disorders |
| 2 | Amla (*Phyllanthus emblica*) | Phyllanthaceae | Leaves & Fruit | Immunity Booster, Antioxidant, Digestive Health |
| 3 | Beil (*Aegle marmelos*) | Rutaceae | Leaves | Digestive Aid, Anti-inflammatory, Respiratory Health |
| 4 | Moringa (*Moringa oleifera*) | Moringaceae | Leaves | Nutrient-Rich, Anti-inflammatory, Antioxidant |
| 5 | Lemon (*Citrus limon*) | Rutaceae | Fruit | Detoxification, Digestive Health, Skin Care |
| 6 | Garlic (*Allium sativum*) | Amaryllidaceae | Bulb | Antibacterial, Antiviral, Heart Health |
| 7 | Ginger (*Zingiber officinale*) | Zingiberaceae | Rhizome | Digestive Aid, Anti-inflammatory, Nausea Relief |
| 8 | Basil (*Ocimum basilicum*) | Lamiaceae | Leaves | Antioxidant, Anti-inflammatory, Digestive Health |

**Table 2:** Inhibition zone (mm) (Mean ± SD) extract of herbsagainst *S. moniliformis*.

| Herbal Extract | Inhibition zone (mm) (Mean ± SD) |
| --- | --- |
| Neem (*Azadirachta indica*) | 14.5 ± 1.22 |
| Amla (*Phyllanthus emblica*) Leaf & Fruit | 22.7±1.44 & 23.5±1.22 |
| Beil (*Aegle marmelos*) | 28.17±1.44 |
| Moringa (*Moringa oleifera*) | 17.0±0.62 |
| Lemon (*Citrus limon*) | 27.0±1.63 |
| Garlic (*Allium sativum*) | 23.2±1.13 |
| Ginger (*Zingiber officinale*) | 21.02±1.05 |
| Basil (*Ocimum basilicum*) | 26.33±1.24 |

Fresh and high-quality plant materials were collected from local sources, ensuring they were free from chemical contaminants or pesticides [18]. Each sample was carefully selected based on morphological characteristics to ensure quality and consistency. To verify species identity, a botanist authenticated each plant specimen [19]. The collected plant materials were then washed thoroughly with distilled water to remove any surface impurities, followed by air-drying under shade at room temperature to preserve their bioactive compounds [20]. Once dried, the plant materials were ground into fine powder using a mechanical grinder and stored in airtight containers until further processing [21].

he selection of these herbal extracts was based on their known phytochemical compositions, which include flavonoids, alkaloids, tannins, essential oils, and other bioactive compounds that have been reported to exhibit antimicrobial activity [22]. These compounds have been studied for their potential to disrupt bacterial cell walls, inhibit enzyme activity, and interfere with quorum sensing mechanisms [23]. By incorporating these plant extracts, the study aims to explore their effectiveness as natural alternatives to conventional antibiotics against *Streptobacillus moniliformis* [24].

### 2.2. Sample Preparation

The collected plant materials were first thoroughly washed with tap water to eliminate any dirt, dust, or potential contaminants [25]. After washing, they were air-dried at **37°C (room temperature)** to ensure moisture removal while preserving their bioactive compounds [26]. To further purify the materials, they were rinsed with **70% ethanol** to eliminate any microbial presence before being washed again with distilled water [27]. Once dried completely, the plant materials were ground into a fine, uniform powder using a **mechanical grinder** [28]. This grinding step was essential to maximize the **surface area for extraction**, ensuring efficient penetration of solvents during the extraction process [29].

### 2.3. Extraction Process

For extraction, **five grams** of each powdered plant sample were weighed and immersed in **50 milliliters of ethanol** [30]. The mixture was left to stand for **72 hours** to allow the active comsspounds to dissolve efficiently [31]. To improve the extraction process, the mixtures were periodically shaken to ensure proper interaction between the plant material and the solvent. After **72 hours**, the extracts were filtered using **Whatman No. 1 filter paper** to remove solid residues [32]. The filtrates were then concentrated at **40°C using a rotary evaporator** to remove excess ethanol, and the crude extracts were stored at **4°C** until further analysis [33].

### 2.4. Bacterial Strain

The bacterial strain used in this study was *Streptobacillus moniliformis*, chosen due to its clinical relevance and limited survival period [34]. The strain was obtained from a **microbiological laboratory** and cultivated on **Mueller-Hinton Agar (MHA) plates** to ensure optimal growth conditions [35]. A **single bacterial colony** was carefully selected from the MHA plate and inoculated into **10 milliliters of Mueller-Hinton Broth (MHB)** to establish an active bacterial culture [36]. The culture was incubated at **37°C for 24 hours** to reach the desired bacterial concentration before being used in antimicrobial testing [37]. Fresh **MHA plates** were prepared for the agar well diffusion method [38].

### 2.5. Agar Well Diffusion Method

The **agar well diffusion method** was utilized to evaluate the antimicrobial activity of the plant extracts against *S. moniliformis* [39]. Mueller-Hinton Agar plates were prepared and solidified under sterile conditions. A **sterile cork borer** was used to create **five wells per plate**, each measuring **6 mm in diameter and 4 mm in depth**, with a **3 cm gap** between wells [40]. **100 microliters** of each herbal extract were introduced into their respective wells. The plates were incubated at **37°C for 24 hours**, and inhibition zones were measured. **DMSO** served as the negative control, ensuring that inhibition was due to the extracts, not the solvent [41].

### 2.6. Incubation and Measurement

After inoculation with the bacterial culture, the prepared plates were incubated at **37°C for 24 hours** under controlled conditions [42]. Following incubation, the zones of inhibition—clear regions indicating bacterial growth suppression—were measured using a **vernier caliper or a millimeter ruler** [43]. These inhibition zones provided a quantitative measure of the **antibacterial activity** of each plant extract. The diameters of the zones were recorded carefully, ensuring accuracy in data collection [44].

Comparisons were made between different extracts to identify the most effective herbal treatments against *S. moniliformis*. Controls were also measured to verify the validity of the results [45].

### 2.7. Quantitative Analysis

All experiments were performed in **triplicate** to ensure the reliability and reproducibility of the results [46]. The data collected were expressed as **mean ± standard deviation (SD)** to account for any variations between replicates [47]. A **p-value of <0.05** was considered statistically significant, indicating that the observed antimicrobial activity was not due to chance [48]. This statistical analysis validated the effectiveness of the plant extracts and helped in identifying the most promising candidates for further antimicrobial applications [49].

## 3. Result

The antibacterial activity of various herbal extracts against *Streptobacillus moniliformis* was assessed using the **agar well diffusion method** [50]. Among the tested extracts, **Beil leaves** and **Lemon peels** demonstrated the most significant antimicrobial effects, producing clear zones of inhibition. Beil leaves exhibited an inhibition zone of **23.5 ± 1.22 mm**, while Lemon peels showed an inhibition zone of **23.27 ± 1.44 mm**. In contrast, **Neem leaves and Moringa extracts** did not display any inhibitory effect, indicating no antimicrobial activity against *S. moniliformis*. These results suggest that certain herbal extracts possess bioactive compounds capable of suppressing bacterial growth, while others lack significant antibacterial properties [51].

**Fig. 1:**
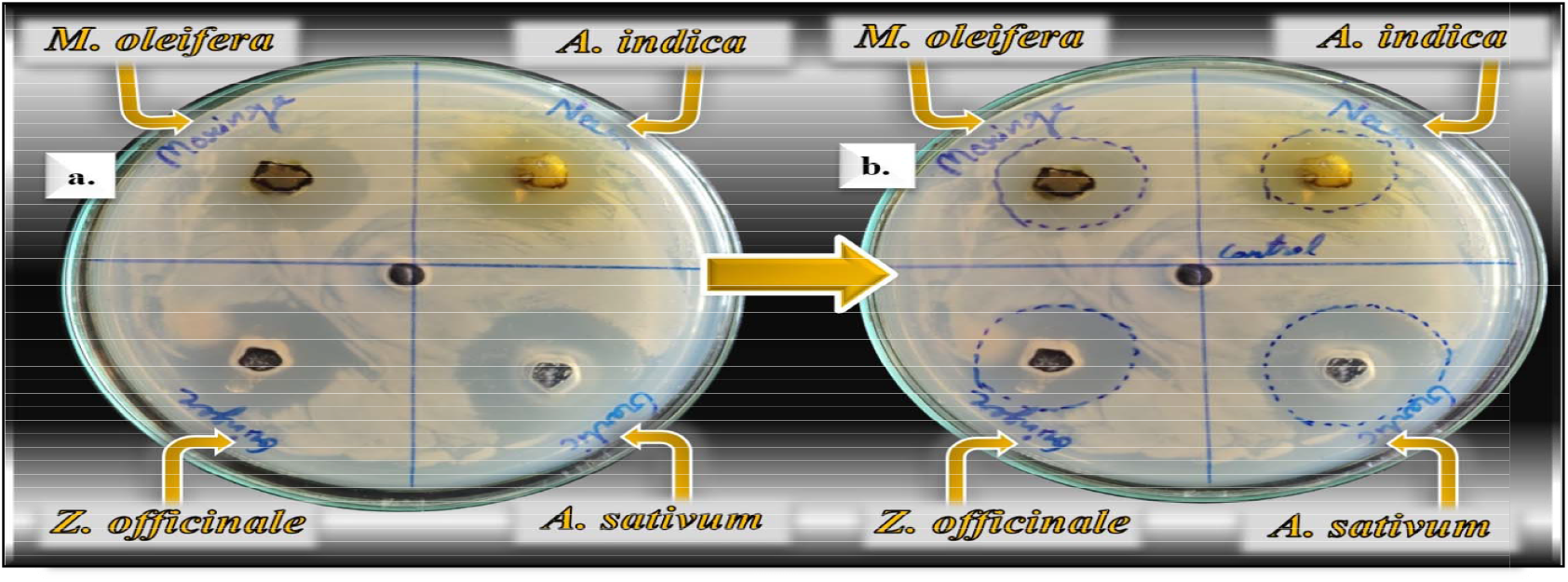
Growth inhibition of multidrug-resistant (MDR) Streptobacillus moniliformis by herbal extracts (i.e. Neem, Moringa, Ginger & Garlic); Without marking ZOI (a) & marking ZOI (b)

**Fig. 2:**
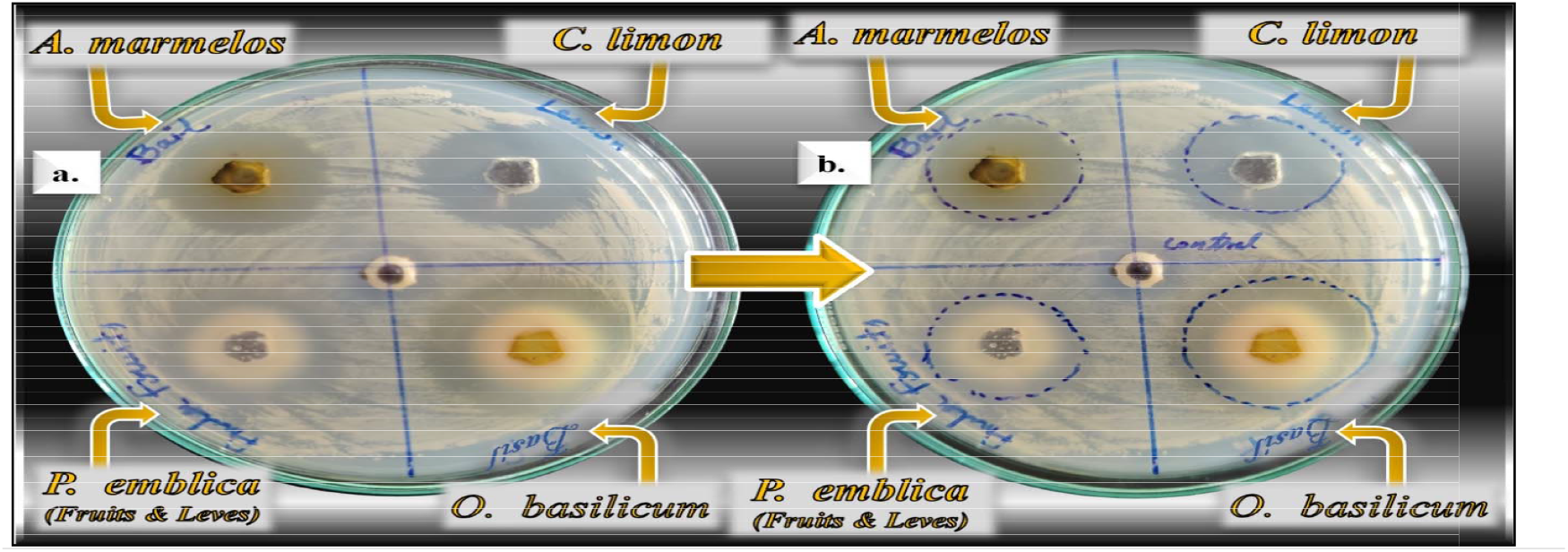
Growth inhibition of multidrug-resistant (MDR) Streptobacillus moniliformis by herbal extracts (i.e. Bael, Lemon, Basi l, & Amla); Without marking ZOI (a) & marking ZOI (b)

### 3.1. Antibacterial potential of different herbal extracts against *S. moniliformis*

The antibacterial potential of various herbal extracts against *Streptobacillus moniliformis* was assessed using the **agar well diffusion method**, with the results reported as **mean ± standard deviation (SD)**. The extracts were further evaluated for their **minimum inhibitory concentration (MIC)** and **minimum bactericidal concentration (MBC)** to determine their antibacterial efficacy [52]. Among the herbal extracts, **Bael leaves (Aegle marmelos)** and **Lemon peels (Citrus limon)** showed the highest antibacterial activity, producing notable zones of inhibition, which suggest their potent antibacterial effects against *S. moniliformis*. **Beil leaves** exhibited an inhibition zone of **23.5 ± 1.22 mm**, while **Lemon peels** produced an inhibition zone of **23.17 ± 1.44 mm** [53].

On the other hand, **Neem leaves (Azadirachta indica)** and **Moringa extracts (Moringa oleifera)** showed no significant inhibition against *S. moniliformis*, with minimal zones of inhibition [54]. Other herbal extracts such as **Ginger peels (Zingiber officinale), Basil (Ocimum basilicum)**, and **Adarak peels** exhibited limited antibacterial activity, with inhibition zones ranging from **14.0 ± 0.82 mm** to **17.33 ± 1.25 mm. Garlic (Allium sativum)** displayed no antibacterial effect, with a zone of inhibition of **0.00 ± 0.00 mm**, likely due to the instability of its bioactive compound **allicin** during the extraction process [55].

Statistical analysis revealed significant differences (**p < 0.05**) in the antibacterial activities among the extracts. **Tukey’s post-hoc test** indicated that **Bael leaves** and **Lemon peels** were significantly more effective than other extracts, such as **Moringa** and **Basil**. These findings underscore the strong antibacterial potential of **Bael leaves** and **Lemon peels**, suggesting that these extracts could be valuable candidates for the development of natural antimicrobial agents [56].

**Fig. 3:**
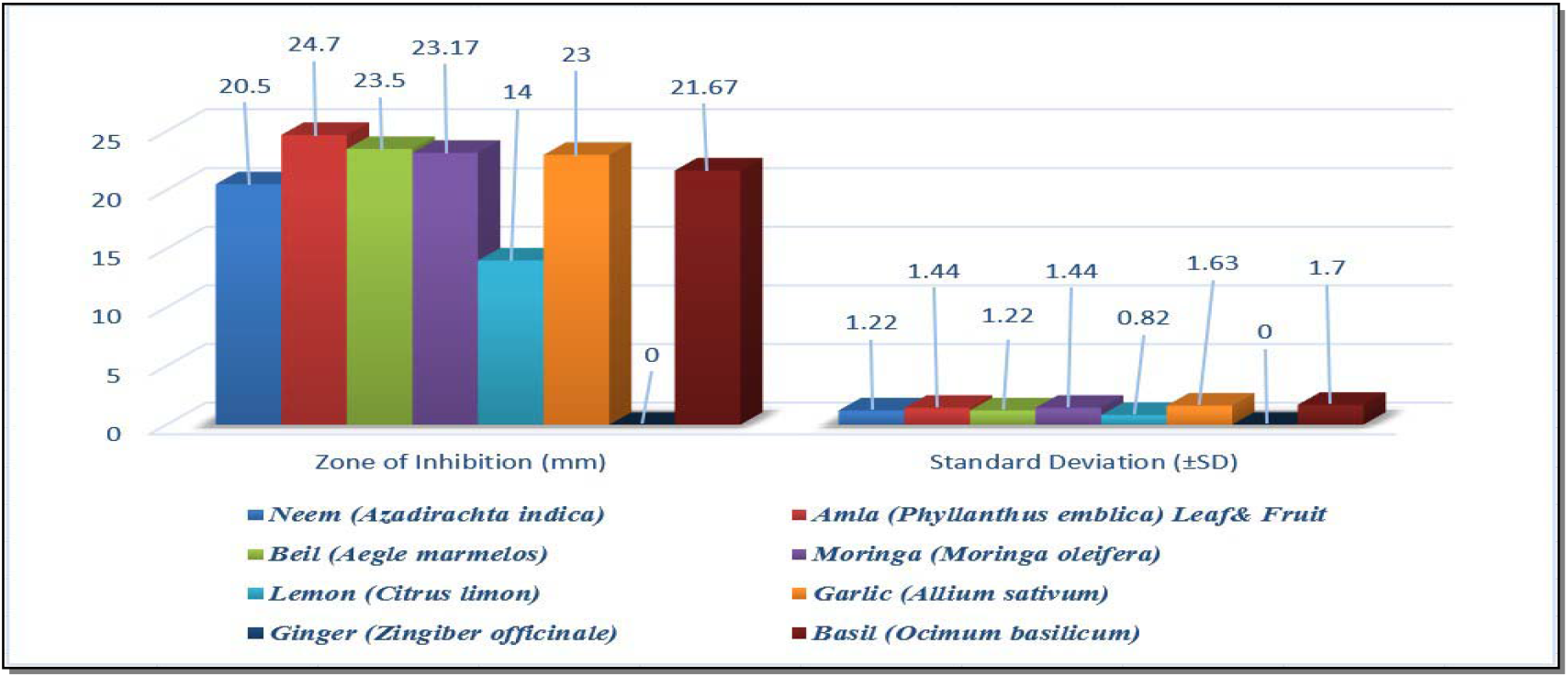
Evaluation of the antibacterial properties extract of herbs against *S. moniliformis*.

The results emphasize the importance of isolating and characterizing the specific bioactive compounds responsible for the antibacterial activity, particularly in **Beil** and **Lemon** extracts, as they show the highest efficacy against *S. moniliformis* [57]. This study serves as a stepping stone toward identifying alternative herbal therapies to combat bacterial infections and address the growing issue of antibiotic resistance [58].

## Discussion

A The rise in multidrug-resistant (MDR) pathogens, particularly *Pseudomonas aeruginosa*, presents a growing global health challenge. This opportunistic bacterium is responsible for severe infections, especially in immunocompromised individuals, highlighting the urgent need for alternative antibacterial treatments. In this context, the present study explored the antibacterial efficacy of twelve herbal extracts against *P. aeruginosa* using the agar well diffusion method. The results reveal that several plant extracts, including **Clove (Syzygium aromaticum), Amla (Phyllanthus emblica)**, and **Lemon (Citrus limon)**, exhibited significant antibacterial activity, showing promise for combating MDR infections. Clove extract demonstrated the highest antibacterial efficacy, with the largest zone of inhibition (**24.33 ± 2.05 mm**), consistent with previous studies that attribute its potent antibacterial effects to **eugenol**, a phenolic compound. Eugenol disrupts bacterial cell membranes, inhibits biofilm formation, and interferes with bacterial enzyme systems, making clove a strong candidate for use in therapeutic applications, especially as a natural alternative to synthetic antibiotics.

Amla leaf and fruit extracts also showed significant antibacterial potential, with inhibition zones of **24.7 ± 1.44 mm** and **23.5 ± 1.22 mm**, respectively. The antibacterial properties of amla can be attributed to its diverse phytochemical content, including **tannins, flavonoids, quercetin**, and **vitamin C**, which interfere with bacterial quorum sensing, destabilize cell walls, and inhibit biofilm formation. This dual action of the fruit and leaf extracts highlights amla’s versatility as a potent antibacterial agent. Similarly, **Cinnamon** and **Lemon** extracts demonstrated noteworthy antibacterial activity, with inhibition zones of **23.0 ± 1.63 mm** for both. The **cinnamaldehyde** in cinnamon interferes with bacterial membrane integrity and metabolic processes, while lemon’s antibacterial activity is attributed to **citric acid, flavonoids**, and **vitamin C**, which alter pH levels and damage bacterial membranes. Both have been previously shown to exhibit broad-spectrum antimicrobial activity, including against *P. aeruginosa*, further supporting their potential as natural antibacterial agents.

**Neem (Azadirachta indica)** and **Thyme (Thymus vulgaris)** also exhibited moderate antibacterial activity, with inhibition zones of **20.5 ± 1.22 mm** and **21.67 ± 1.7 mm**, respectively. The antibacterial effects of neem can be attributed to **azadirachtin** and other **limonoids**, which prevent bacterial growth and reduce pathogenicity, while thyme’s antibacterial properties are linked to **thymol** and **carvacrol**, which disrupt bacterial membrane integrity and respiration. While their antibacterial effects were moderate, these plants could serve as supplementary agents in combination therapies to enhance the overall effectiveness of antimicrobials.

Interestingly, extracts of **Basil (Ocimum basilicum), Moringa (Moringa oleifera)**, and **Ginger (Zingiber officinale)** displayed weaker antibacterial activity, with inhibition zones of **13.33 ± 1.24 mm, 17.33 ± 1.25 mm**, and **14.0 ± 0.82 mm**, respectively.

The low antibacterial efficacy of ginger could be attributed to the instability of **gingerol**, its key bioactive compound, during extraction or storage. Similarly, the reduced activity of basil and moringa may be due to lower concentrations of bioactive compounds or reduced efficacy against *P. aeruginosa*. Additionally, **Garlic (Allium sativum)**, known for its antimicrobial compound **allicin**, showed no antibacterial activity, likely due to allicin’s degradation under the extraction conditions.

Statistical analysis using different test revealed significant differences (**p < 0.05**) in the antibacterial efficacy of the extracts. **Amla**, and **Lemon** extracts were found to be highly effective compared to the weaker extracts like **Moringa** and **Basil**, emphasizing the potential of these plants as natural antibacterial agents. The phytochemical profiles of these extracts suggest that they target multiple bacterial pathways, which may reduce the risk of resistance development—a crucial advantage over synthetic antibiotics that often target specific bacterial structures.

The findings underscore the potential of herbal extracts as **sustainable** and **eco-friendly** alternatives to synthetic antibiotics. These extracts, with their broad-spectrum activity and lower likelihood of resistance development, could contribute significantly to combating the global issue of antibiotic resistance. Furthermore, their biodegradability, low toxicity, and alignment with sustainable healthcare practices make them promising candidates for **integrative therapeutic strategies**. As the incidence of MDR infections continues to rise, these herbal extracts could play an important role in future antibacterial treatments, offering a natural, effective, and sustainable approach to infection management.

## Conclusion

The findings of this study demonstrate that herbal extracts, particularly **Beil (Aegle marmelos), Amla (Phyllanthus emblica)** and **Lemon (Citrus limon)**, possess significant antibacterial activity against the multidrug-resistant pathogen **Pseudomonas aeruginosa**. The inhibitory effects observed can be attributed to potent bioactive compounds such as **eugenol, tannins, flavonoids**, and **citric acid**. These compounds disrupt bacterial cell membranes, inhibit quorum sensing, and prevent biofilm formation, thereby inhibiting bacterial growth. The study underscores the potential of these natural products as environmentally friendly and sustainable alternatives to synthetic antibiotics. Future research should focus on isolating and characterizing these bioactive compounds to develop effective therapeutic agents for treating antibiotic-resistant bacterial infections.

## Acknowledgments

We would like to express our heartfelt gratitude to **Shobhit Institute of Engineering & Technology, Meerut**, and **Shobhit University, Gangoh**, for providing the necessary tools and resources that facilitated the completion of this study. Our sincere thanks also go to the **School of Biotechnology & Life Sciences** for their unwavering support and guidance throughout this research. We are deeply grateful to our lab staff and colleagues for their assistance with sample preparation, analysis, and data collection. Their contribution was indispensable to the success of this work.

We also wish to acknowledge the valuable contributions of the **microbiologists** and **botanists** whose expertise helped confirm the identification of both bacterial strains and plant species. Their input played a critical role in ensuring the accuracy and reliability of our results.

Lastly, we would like to extend our heartfelt appreciation to our peers and families for their constant encouragement and support. Without their collaborative efforts and dedication, this study would not have been possible.

## Statements and Declarations

The research described in this publication was conducted without any conflicting financial or non-financial interests influencing the work. There was no external funding or support received for this study. Ethical approval was not required for the research.

The authors’ contributions are as follows: **Kishlay Kant Singh** (first author) was responsible for the overall conceptualization, methodology, data collection, and manuscript writing. **Divya Prakash** (guide) provided mentorship, supervision, and guidance throughout the research process. **Mansi Saini** contributed significantly to the drafting, revising, and refinement of the manuscript. The experiments were performed by the authors and laboratory staff, who also assisted in gathering data.

The corresponding author will provide the data supporting the conclusions of this study upon reasonable request. Since the research did not involve human participants, consent for participation was not applicable. All authors have read, reviewed, and approved the final manuscript for submission and publication.

